# Single cell mass spectrometry reveals intercellular compartmentalization of camptothecin biosynthesis in the tree *Camptotheca acuminata*

**DOI:** 10.1101/2025.06.24.661346

**Authors:** Van-Hung Bui, Anh Hai Vu, C. Robin Buell, Chenxin Li, Lorenzo Caputi, Thu-Thuy T. Dang

## Abstract

The medicinal tree *Camptotheca acuminata* produces camptothecin, a monoterpenoid indole alkaloid (MIA) precursor for several leading chemotherapeutic agents (Lorence and Nessler 2004). Although the biosynthesis of camptothecin remains poorly understood, a putative route has been hypothesized based on *in planta* metabolite profiling and feeding studies (Fig. 1a) (Sheriha and Rapoport 1976; Sadre et al. 2016). However, pathways proposed on whole-tissue or organ-level metabolomic and transcriptomic data lack resolution on the intricate, cell-specific compartmentalization of natural products biosynthesis. Such gaps could be addressed by single cell technologies, which have recently shown tremendous potential to transform gene discovery in herbaceous plants (Li et al. 2023; Zhan et al. 2023; Vu et al. 2024; Wu et al. 2024; McClune et al. 2025). Nevertheless, single cell mass spectrometry (scMS) has not been adapted for woody species, largely due to the challenges associated with their highly lignified tissue and complex cellular architecture. In this study, we developed an scMS pipeline for the woody tree *C. acuminata* to investigate the intercellular organization of camptothecin biosynthesis.

---

Prior to single cell analyses, we conducted isotope feeding experiments using *d5*-tryptamine (Fig. S1) to identify sites of active camptothecin biosynthesis. All tested tissues showed substantial amounts of labeled MIAs, especially *d4*-strictosidinic acid, *d4*-strictosamide, and *d4*-vincosamide, confirming that early-stage biosynthetic activity is widespread. In contrast, roots accumulated only trace amounts of labeled intermediates, indicating limited biosynthetic activity. Notably, only young and mature stems produced detectable amounts of labeled camptothecin, suggesting that the stem is the primary site of camptothecin biosynthesis in *C. acuminata*. To investigate the cell-type compartmentalization of camptothecin biosynthesis, we first generated protoplasts using protocols optimized for shoot apical meristem, leaf, and stem of *C. acuminata* (Li et al. 2023; Vu et al. 2024) (Fig. 1b). The resulting protoplasts were sorted into 96-well plates using an automated single cell picking system. Following the addition of solvent and internal standard, each cell was analyzed by ultra-high-performance liquid chromatography coupled with high resolution mass spectrometry (UHPLC-HRMS). A total of 576 single cells per tissue type were initially collected across six 96-well plates. To minimize batch effects from technical variability, each plate was treated as an independent experiment. After excluding wells that were empty or contained more than one cell, the final dataset included 495 shoot apical meristem cells, 495 leaf cells, and 493 cells from the stem (Supplementary Datasets 1–3). Targeted metabolites were quantified based on external calibration with authentic standards (Tables S1 and S2), and analyte concentrations were normalized to the cell volume, calculated based on the diameters measured during single cell collection.

**Fig. 1.**
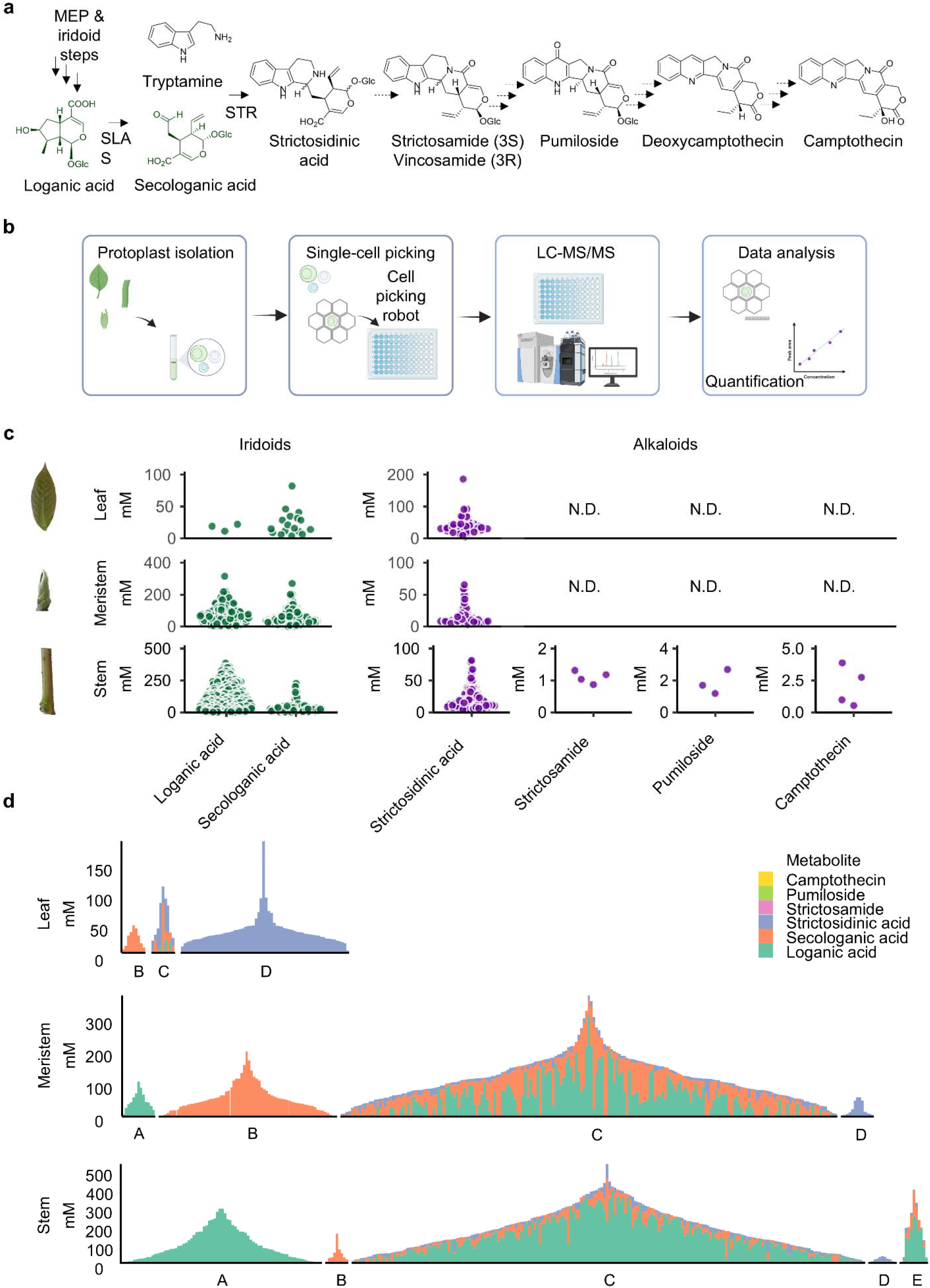
Single cell metabolomics of *Camptotheca acuminata.* (a) Proposed camptothecin biosynthetic pathway. (b) Schematic experimental pipeline. (c) Dot plots showing concentrations of key metabolites in single cells across organs. Each data point represents a protoplast. For each metabolite, dots were offset from the center based on the distribution of concentrations among single cells. (N.D.: not detected). (d) Stacked bars showing concentrations in individual cells of loganic acid, secologanic acid, strictosidinic acid, strictosamide, pumiloside, and camptothecin). Cells are organized into groups with loganic acid only (A), secologanic acid only (B), loganic and secologanic (C), strictosidinic acid only (D), and other alkaloids (strictosamide, pumiloside, and camptothecin) (E). In each group, cells are ordered by total metabolite concentration with highest concentration in the center and lower producers closer to the edges. Panel widths scale with the number of cells in each group, and colours cor- respond to individual metabolites.

Single cell profiling revealed that major iridoids accumulated at remarkably high intracellular concentrations, often reaching the millimolar range (Fig. 1c). Among these, loganic acid levels were highest in stem (128 mM), followed by meristem (74 mM) and leaf (17 mM) cells. Notably, cells accumulating loganic acid comprised a substantial fraction of the cells obtained from stem (56.6%) and meristem (31.7%) but were rare in leaves (0.6%). Secologanic acid, the immediate downstream product of loganic acid, also reached millimolar concentrations across all cells. Strictosidinic acid, a key intermediate in camptothecin biosynthesis, was detected across all tissues, with the highest prevalence in meristem (52.9%), followed by stem (38.3%), and very low abundance in leaf (3.4%) cells. In contrast, strictosamide, pumiloside, and camptothecin were exclusively detected in 1% of the total stem protoplasts, suggesting that camptothecin biosynthesis is restricted to a rare and specialized cell type within the stem. Co-occurrence analysis of metabolites in stem tissues identified five distinct cells populations: cells containing only loganic acid (A); only secologanic acid (B); both loganic acid and secologanic acid (with or without strictosidinic acid) (C); only strictosidinic acid (D); and cells containing camptothecin and its proposed intermediates (strictosamide, pumiloside, camptothecin) (E) (Fig. 1d). Importantly, most cells accumulating loganic acid also contained secologanic acid across all tissues. However, co-occurrence of early-stage iridoid intermediates (loganic acid, secologanic acid) with late-stage camptothecin intermediates was rare, reinforcing the notion that late stage camptothecin biosynthesis occurs in a distinct, specialized population. These findings highlight a cellular compartmentalization of the pathway, with early iridoid biosynthesis localized to one cell type and downstream alkaloid biosynthesis restricted to another rare population. Interestingly, the scMS data revealed that the metabolite distributions observed in bulk tissue analyses are driven by a small number of MIA-rich cells, rather than being evenly distributed throughout the tissue. Comparative scMS analysis across tissues also confirmed that neither leaf nor meristem contained detectable levels of camptothecin. This spatial separation of biosynthetic activity likely reflects differences in transcriptional regulation across organs, possibly driven by organ-specific expression of transcription factors.

scMS of arboraceous species like *C. acuminata* presents several technical challenges, including lignified secondary cell walls, low abundance and instability of pathway intermediates, potential organelle-level metabolite compartmentalization, and the lack of well-established cell-type markers. Nevertheless, the cell-level spatial compartmentalization of camptothecin and its precursors demonstrated here could be combined with single cell transcriptomic approaches to ultimately enable the elucidation of the camptothecin biosynthetic pathway. Overall, these findings demonstrate the power of scMS to uncover the biosynthesis and intercellular transport of specialized metabolites in woody species such as *C. acuminata*.

## Acknowledgements

CRB receives funding from Georgia Research Alliance, Georgia Seed Development, University of Georgia. C.R.B and C.L. acknowledge funding from the National Science Foundation MCB-2309665 (C.R.B and C.L.). LC is grateful for the Max Planck Gesellschaft. T.T.T.D. receives funding from NSERC Alliance Collaboration (ALLRP 571673 – 21) and Catalyst (ALLRP 579871 – 22), and the Michael Smith Health BC Scholar Award (SCH-2020-0401). We thank Dr. Maritta Kunert and Sarah Heinicke for their assistance with MS analyses, Jens Wurlitzer for the cell-picking robot, Eva Rothe and the greenhouse team for plant care, and Dr. Moonyoung Kang and Tingan Zhou for their advice on the protoplast isolation protocol.

## Author contributions

CL, C.R.B., LC and T.T.T.D. designed the study. V.H.B. generated single cell metabolomics data with assistance from A.H.V. and L.C. C.L. and V.H.B. performed data analyses. C.L. and V.H.B wrote the manuscript with input from all authors.

## Data Availability

Individual cell level metabolite abundances are available in Supplementary Datasets 1–3.

## Supporting Information

1) Supplemental methods. 2) Supplementary Table S1-S2. 3)-5) Supplementary Datasets 1–3.

