## Supplemental data for "Single cell mass spectrometry reveals intercellular compartmentalization of camptothecin biosynthesis in the tree *Camptotheca acuminata*"

**Table of contents**

**Experimental procedures**	S3-5

**Supplementary Figures**

**Supplementary Tables**

**Experimental procedures**

Methods S1: Chemicals

Information about chemicals, reagents, and solvents is listed in Table S1.

Methods S2: Plant growth conditions.

One-year-old *C. acuminata* plants were obtained from Planten Tuin Esveld, Netherlands. Plants were cultivated in a standard soil mix in a greenhouse at 20-25°C (daytime) and 17-20°C (nighttime) with 40-70% humidity under the minimum artificial light of 12 hrs at Jena Germany.

Methods S3: Feeding of *C. acuminata* tissue

For isotopologue feeding experiments, fresh plant tissues (10-mm leaf disks, 5-mm meristem segments, 5-mm stem segments, and 5-mm root segments) were excised and incubated individually in 200 µL of aqueous solution containing 500 μM d5-tryptamine for 5 days in a growth chamber under conditions of 16-hour daylight at 23°C, followed by night conditions at 21°C, with humidity levels maintained between 40% and 60%. After incubation, residual liquid was discarded, and metabolites were extracted from the tissues using extraction solvent (85% methanol, 15% water, 0.1% formic acid). The extracts were filtered through 0.22 µm PTFE syringe filters. Any remaining liquid was removed, and the samples were extracted with extraction solvent (85% methanol, 15% water, 0.1% formic acid), filtered, and then diluted 10-fold with extraction buffer containing 50 nM catharanthine as an internal standard before LC-MS.

Methods S4: Leaf and meristem protoplast isolation for single cell metabolomics.

Young leaves (100 mg) measuring ~3 cm in length from the first and second nodes were selected, rinsed gently with water, and cut into 1 mm strips using a sterile surgical blade. Leaf strips were immediately transferred to a Petri dish containing 10 mL of digestion medium composed of 2% (w/v) Cellulase Onozuka R-10, 0.3% (w/v) Macerozyme R-10, and 0.1% (v/v) pectinase in Mannitol-MES (MM) buffer (0.4 M mannitol, 20 mM MES, pH 5.7–5.8 adjusted with 1 M KOH). The open Petri dish was placed in a desiccator and subjected to a vacuum of 100 mBar for 15 min, with periodic vacuum disruption (10s per minute), facilitating infiltration of the digestion medium into the leaf tissue. Following vacuum infiltration, the leaf strips were incubated in the digestion medium at room temperature for 2.5 hours. Post-incubation, the mixture was gently agitated on an orbital shaker at approximately 70 rpm for 30 minutes to facilitate protoplast release. To remove cellular debris, the protoplast suspension was filtered sequentially through nylon sieves of 70 µm and 40 µm mesh sizes. The filtrate was centrifuged at 200×g for 5 minutes at 4°C with gentle acceleration and deceleration to pellet the protoplasts. The supernatant was removed, and the protoplast pellet was washed once with 5 mL of MM buffer. Protoplasts were then pooled and resuspended in 2 mL of MM buffer. For protoplast purification via OptiPrep density gradient, a working solution (WS) was prepared by dissolving 6 mg KCl in 1 mL OptiPrep. Layer A was created by mixing 500 µL WS with 2 mL protoplast suspension. Layer B consisted of 266 µL WS combined with 1 mL resting solution, and Layer C contained 200 µL MM buffer alone. Layers were sequentially and carefully layered into a 5 mL tube (A, followed by B and C) without disturbing interfaces. The gradient was centrifuged at 200×g for 5 minutes at 4°C, allowing viable protoplasts to float to the top and debris to settle at the bottom. The uppermost 200 µL fraction containing purified protoplasts was collected into a new tube. Protoplast viability and concentration were determined using fluorescein diacetate staining and hemocytometer counting. The final concentration was adjusted to 1 × 10⁶ protoplasts per mL.

Methods S5: Stem protoplast isolation for single cell metabolomics and single cell RNA-seq.

Young stems from the 2nd to 4th nodes below the shoot apex were sampled in ice-cold water in a 50 mL tube. Using a sterile razor blade, an incision was made longitudinally along the stem segments, and the outer stem layers were carefully peeled away. Both outer and inner stem tissues were immediately transferred into 20 mL of protoplast enzyme cocktail. The cocktail contained 0.4 M sorbitol, 20 mM KCl, 20 mM MES, 2% (w/v) Cellulase Onozuka R-10, 0.3% (w/v) Macerozyme R-10, 0.1% (v/v) pectinase, 1 mM CaCl₂, 0.2% (v/v) 2-mercaptoethanol, and 0.1% (w/v) bovine serum albumin (BSA), adjusted to pH 5.7 and heated at 50°C for 10 minutes before cooling to room temperature. Vacuum infiltration of stem segments in the protoplast cocktail was performed for 10 minutes. Following vacuum infiltration, the mixture was shaken at 80 rpm for 70 minutes at room temperature. Subsequently, the mixture was briefly shaken at 100 rpm for 5 minutes to facilitate protoplast release. The suspension was then filtered through sequential nylon sieves of 100 µm and 70 µm mesh sizes to remove debris. Filtrate was centrifuged at 200 × g for 5 minutes at 4°C, after which the protoplast pellet was gently resuspended in 2 mL of resting solution. For protoplast purification via OptiPrep density gradient, the procedure was the same as the leaf/meristem protoplast isolation. Protoplast viability and concentration were determined using fluorescein diacetate staining and hemocytometer counting. The final protoplast concentration was adjusted to 1 × 10⁶ protoplasts per mL.

Methods S6: Single cell picking for metabolite profiling.

This procedure was modified based on the method described by Vu et al. (Vu et al., 2024). Single-cell isolation was performed using SIEVEWELL™ chips (Sartorius) containing 90,000 nanowells (50 μm diameter × 50 μm depth). The chips were initially primed with 100% ethanol, rinsed twice with MM buffer, and incubated with 5% BSA in MM buffer for 30 min at room temperature. After removing the BSA solution through side ports, chips were replenished with fresh MM buffer. Approximately 10,000 diluted protoplasts suspended in 1 mL of MM buffer were then gently dispensed across each chip in a Z-shaped pattern. Excess solution was drained via the side ports. Chips were subsequently mounted on a CellCelector™ Flex (Sartorius) equipped with an optical imaging system comprising a fluorescence microscope (Spectra X Lumencor) and a CCD camera (XM-10). Nanowells were imaged under bright-field or fluorescence (DAPI filter) illumination. Single protoplasts, along with 20 nL of nanowell solution, were individually collected using a 30 μm glass capillary and transferred into SureSTART™ WebSeal™ 96-well microtiter plates (Thermo Fisher Scientific) containing 8 μL MilliQ water with 0.1% formic acid. Images confirming successful picking were captured before and after cell collection. Following single-cell collection, 8 μL methanol containing 20 nM cathanrantine as an internal standard was added to each well. Pooled quality-control (QC) samples, prepared by combining 2 μL from each sample, were included in each experimental batch for QC purposes and MS² fragmentation analyses.

Methods S7: LC-MS analysis.

UHPLC-HRMS analysis was conducted using a Vanquish UHPLC system coupled to a Q-Exactive Plus Orbitrap mass spectrometer (Thermo Fisher Scientific). Metabolites were separated on a Waters™ ACQUITY UPLC BEH C18 130 Å column (1.7 μm particle size, 1 mm × 50 mm) maintained at 40 °C. The mobile phase consisted of a binary gradient system with solvent A (MilliQ water with 0.1% formic acid) and solvent B (acetonitrile, ACN). The gradient started at 1% solvent B and increased linearly to 70% solvent B over 5 min, followed by a washing step at 99% solvent B for 0.5 min. The column was subsequently re-equilibrated at 1% solvent B for 1.5 min, resulting in a total run time of 7 min. The flow rate was maintained at 0.3 mL min⁻¹ with an injection volume of 4 μL. During the analysis, samples were held at 10 °C in the autosampler, and the autosampler needle was cleaned for 20 s after each injection using a 1:1 (v/v) mixture of methanol and MilliQ water at a washing speed of 50 μL s⁻¹. The Q-Exactive Plus Orbitrap mass spectrometer was operated with a heated electrospray ionization (HESI) source, calibrated using Pierce positive and negative ion mass calibration solutions (Thermo Fisher Scientific). The following HESI parameters were applied based on the UHPLC flow rate of 0.3 mL min⁻¹ in source auto-default mode: sheath gas flow rate of 48, auxiliary gas flow rate of 11, sweep gas flow rate of 2, spray voltage of +3,500 V, capillary temperature of 256 °C, auxiliary gas heater temperature of 413 °C, and S-lens RF level of 50. Data acquisition was performed in positive mode using full-scan MS with a resolution of 70,000 (FWHM at 200 Da), scanning across a mass range of *m/z* 120–1,000. Quality control (QC) pooled samples were analyzed using data-dependent MS/MS (full MS/dd-MS², Top10), enabling simultaneous acquisition of precursor and fragmentation spectra. Additionally, an inclusion list-based dd-MS² analysis was conducted on QC samples to confirm fragmentation patterns of targeted precursors. Parameters for dd-MS² analyses included a resolution of 17,500, an isolation window of 0.7 Da, and normalized collision energies (NCE) at three levels: 15%, 30%, and 45%. Spectra were acquired in centroid mode, and all instrument parameters were controlled via Xcalibur software (version 4.3.73.11, Thermo Fisher Scientific). Standard solutions of alkaloids, and iridoids were prepared at approximately 1 mM in methanol, with exact concentrations recorded. UHPLC-MS analyzed serial dilutions down to 0.01 nM to establish the limit of quantification (LOQ) and calibration ranges, with each calibration concentration measured in triplicate. Chromatographic peak areas from extracted ion chromatograms (EIC) were integrated and quantified using Xcalibur Quan Browser software (version 4.3.73.11, Thermo Fisher Scientific). Concentration of key metabolites across single cells were also reported as Supplementary Datasets 3-5, for leaf, meristem, and stem, respectively.

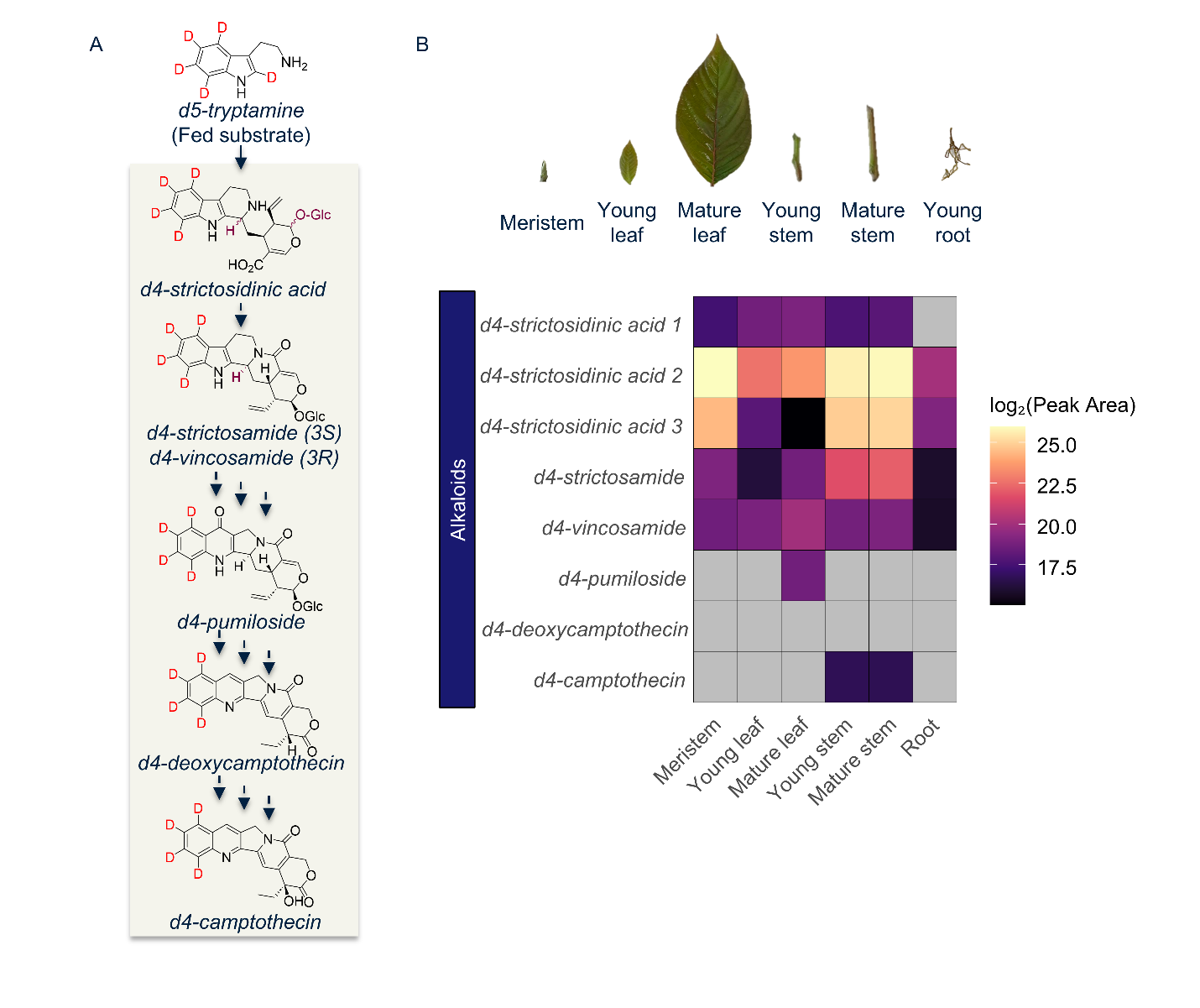

**Figure S1**. Feeding experiments reveal the active tissue(s) of camptothecin biosynthesis.

(A). Structures of labelled substrate and metabolites being monitored.

(B). Heat map showing abundances (log2 peak area) of labelled metabolites across organs. Grey color indicates lack of detection. Photos of organs are shown on the top of the heatmap.

**Table S1.** List of chemicals used in this study.

|  | **Chemical** | **Supplier** |
| --- | --- | --- |
| 1 | Catharanthine | Sigma Aldrich |
| 2 | Loganic acid | Extrasynthese |
| 3 | Secologanic acid | Synthesized in the laboratory |
| 4 | Strictosidinic acid | Synthesized in the laboratory |
| 5 | Strictosamide | Arctom Scientific |
| 6 | Vincosamide | BioBioPha |
| 7 | Pumiloside | BioCrick |
| 8 | Deoxycamptothecin | Synthesized in the laboratory |
| 9 | Camptothecin | Sigma Aldrich |
| 10 | Macerozyme R-10 from Rhizopus sp. Lyophil. | SERVA |
| 11 | Cellulase »Onozuka« R-10 aus Trichoderma viride ca. 1 U/mg | SERVA |
| 12 | Pectinase from Aspergillus niger | Sigma Aldrich |
| 13 | Fluorescein Diacetate | TCI |
| 14 | D-Mannitol | Sigma Aldrich |
| 15 | MES hydrate | Sigma Aldrich |
| 16 | OptiPrep™ Density Gradient Medium | Sigma Aldrich |
| 17 | Bovine serum albumin | Sigma Aldrich |
| 18 | Formic Acid Optima LC/MS | Fisher Scientific |
| 19 | Acetonitrile OPTIMA® LC/MS GRADE | Fisher Scientific |
| 20 | Methanol UHPLC-MS | Fisher Scientific |

**Table S2**. Validation of the targeted scMS analysis.

| **Compound** | **Quantitative Ion (m/z)** | **Calibration Range (nM)** | **Regression**  **Equation ^1^** | **Correlation Coefficient (R2)** | **LOQ ^2^ (nM)** |
| --- | --- | --- | --- | --- | --- |
| Loganic acid | [M+H]+ (337.191) | 2-10000 | y =17184x + 72318 | 0.9999 | 2 |
| Secologanic acid | [M+H]+ (375.128) | 2-10000 | y = 19685x + 47870 | 0.9988 | 2 |
| Strictosidinic acid | [M+H]+ (517.218) | 0.5-1000 | y = 82970x - 59756 | 0.9999 | 0.5 |
| Strictosamide | [M+H]+ (499.207) | 0.5-1000 | y = 116955x + 49869 | 0.9999 | 0.2 |
| Pumiloside | [M+H]+ (513.186) | 1-1000 | y = 25529x - 16957 | 0.9998 | 1 |
| Camptothecin | [M+H]+ (349.118) | 0.5-1000 | y =188859x - 38230 | 0.9999 | 0.2 |

^1^ Each point of calibration curve was measured in triplicate.

^2^ The limit of quantification (LOQ) was defined as the lowest concentration of analyte injected that consistently produced a signal-to-noise (S/N) ratio of ≥10 in triplicate analyses.
